# Fine-scale position effects shape the distribution of inversion breakpoints in *Drosophila melanogaster*

**DOI:** 10.1101/793364

**Authors:** Jakob McBroome, David Liang, Russell Corbett-Detig

## Abstract

Chromosomal inversions are among the primary drivers of genome structure evolution in a wide range of natural populations. While there is an impressive array of theory and empirical analyses that has identified conditions under which inversions can be positively selected, comparatively little data is available on the fitness impacts of these genome structural rearrangements themselves. Because inversion breakpoints can interrupt functional elements and alter chromatin domains, each rearrangement may in itself have strong effects on fitness. Here, we compared the fine-scale distribution of low frequency inversion breakpoints with those of high frequency inversions and inversions that have fixed between *Drosophila* species. We identified important differences that may influence inversion fitness. In particular, proximity to insulator elements, large tandem duplications adjacent to the breakpoints, and minimal impacts on gene coding spans are more prevalent in high frequency and fixed inversions than in rare inversions. The data suggest that natural selection acts both to preserve both genes and larger cis-regulatory networks in the occurrence and spread of rearrangements. These factors may act to limit the availability of high fitness arrangements when suppressed recombination is favorable.

## Introduction

In natural *Drosophila melanogaster* populations many individuals carry inversions, or chromosomes with large genomic regions reversed by large-scale mutations. These rearrangements have a long history of study in *Drosophila* species (Dobzhansky 1962; Sturtevant, 1917). The primary hypothesis explaining the prevalence of inversions in natural populations is that reduced recombination between inverted and standard arrangements in heterozygotes is favored by natural selection (Corbett-Detig & Hartl, 2012; Fuller et al 2019; Kapun et al 2016a; Langley et al 2012; Mukai 1971; Sturtevant and Beadle 1936). Alleles contained in inversions can interact epistatically or additively to maintain a polygenic complex phenotype such as body size, stress resistance, fecundity, and lifespan (Hoffman et al 2004; Hoffman and Rieseberg 2008; Kirkpatrick 2010). If such a complex phenotype is favored by natural selection but broken down by recombination on freely recombining chromosomes, inversions that maintain linkage among genes contributing to that phenotype can be positively selected. Biogeographic data supports this hypothesis; natural populations of *Drosophila melanogaster* maintain inversion frequency clines that are strongly correlated with climatic clines (Kapun et al 2016a; Kapun et al 2016b; Knibb 1982; Mettler et al 1977; Rane et al 2015; Simões and Pascual 2018). Furthermore, evidence consistent with selection on polygenic phenotypes associated with chromosomal inversions is known from an ever expanding set of taxa (Butlin et al 1982; Huynh et al 2011; Oneal et al 2014). It is therefore increasingly widely accepted that a major source of positive selection on chromosomal inversions is the maintenance of linkage among alleles that are favorable in similar contexts.

Whereas the potential fitness effects of decreasing recombination among alleles that are favorable in combination are well established, the impacts of an inversion event on the original individual it occurs in are not well understood. Nonetheless, these impacts are also likely to play an important role in shaping evolutionary outcomes for new arrangements. For example, an inversion breakpoint that interrupts a gene sequence could result in immediate negative fitness effects if that gene is important for an individual’s success. If these kinds of negative fitness effects are prevalent across most of the genome, the rate of formation of inversions which can spread to high frequency is expected to be correspondingly low. Understanding the fitness impacts of chromosomal inversions is fundamental to characterizing their evolutionary potential in natural populations, and therefore requires a deeper characterization of the factors that influence each arrangement’s intrinsic fitness.

Accumulated evidence is consistent with the idea that inversion breakpoint positions might have large impacts on fitness. The distribution of polymorphic inversion breakpoints along the genome is not random (Calvete et al 2012; Gonzalez et al 2007; Orengo et al 2015; Pevzner and Tesler 2003; Puerma et al 2014; Puerma et al 2016; Tonzetich et al 1988). In fact, many independently formed inversions appear to precisely share breakpoint locations (Gonzalez et al 2007; Puerma et al 2014; Pevzner and Tesler 2003). Even when inversion breakpoints are not precisely reused at the molecular level, their broad-scale distribution across the genome is non-uniform (Pevzner and Tesler 2003; Ranz et al 2007). These patterns might be due to mutational bias if these regions are particularly “fragile”, or driven by selection if inversions whose breakpoints occur within specific contexts have higher intrinsic fitness. Therefore, although this pattern is well-established across many species, the factors underlying breakpoint localization and the intrinsic fitness of new arrangements more generally are poorly understood.

We examine two general categories of features that shape the fine-scale distribution of inversion breakpoints. First, mutational biases are factors tied to the likelihood of breakage and inverted repair at a specific genomic location (Calvete et al 2012; Guillen and Ruiz 2012; Pevzner and Tesler 2003; Tonzetich et al 1988). In many species sequence repetition encourage the formation of inversions through ectopic recombination, an example of a mutational bias (Guillen and Ruiz 2012), though this is relatively rare in the *melanogaster* subgroup (Ranz et al. 2007, Corbett-Detig and Hartl 2012). Physical instability due to unstable secondary structure or local chromatin environment are also potential mutational biases that could affect breakpoint localization. Second, it is possible that specific breakpoint positions affect the intrinsic fitness of a new arrangement. We term factors impacting the relative fitness of any given inversion rearrangement selective factors, distinct from mutational biases in that they act after the inversion event to shape the observable distribution of inversion breakpoints. The breakage of an important gene span at the breakpoint would be a negative selective effect, as would the modification of spatial relationships among regulatory elements. These effects define what we term the “intrinsic fitness” of an inversion event, or the fitness impacts of the inversion itself outside of any polygenic trait it confers.

Comparisons among fixed, high frequency and low frequency inversions can reveal the impact of natural selection on chromosomal inversion breakpoints. Because they have persisted within natural populations and risen to high frequencies, high-frequency and fixed chromosomal inversions are expected to show a biased distribution of local breakpoint features consistent with the factors that result in higher fitness arrangements. Conversely, low-frequency inversions are most likely recently arisen arrangements and should primarily reflect mutational biases. Therefore, by examining the distributions of fixed, high frequency and low frequency inversion breakpoints we can identify the factors that shape the intrinsic fitness of newly-arise arrangements.

The primary sequence-level impact of an inversion occurs at its breakpoints, where the genome sequence is interrupted and replaced with an entirely different section of chromosome transposed from a large distance away. The effects of an inversion breakpoint on the genome are sometimes called “position effects” and examples have been identified in a variety of organisms (Casterman et al 2007; Frischer et al 1986; Hough et al 1998; Lakich et al 1993; Puig et al 2004). Position effects include the disruption of important contiguous sections of the genome such as gene coding spans (Frischer et al 1986) and the transformation of the local regulatory landscape by enhancer shuffling (Ren and Dixon 2015) or chromatin relocation (Cryderman et al 1998; Eissenberg et al 1992).

We expect these disruptive position effects to usually have negative impacts on inversion fitness. If severe fitness penalties are imposed for breakage in a large proportion of the genome, correspondingly few random inversions will have a neutral or positive intrinsic fitness. Negative position effects could easily be stronger than any positive impacts from the maintenance of allelic combinations that confer complex phenotypes, limiting the rate of formation of polymorphic inversions. Positively selected position effects are also possible; previous work has shown that breakpoint localization near “sensitive sites” strongly intensifies the often-positive recombination suppression effect (Corbett-Detig 2016; Coyne et al 1993; Hawley 1980; Navarro and Ruiz 1997; Roberts et al 1970). Despite these considerations, breakpoint effects have generally been secondary considerations in the study of *Drosophila* inversion evolution (Puig et al 2004).

We considered several specific candidate factors for selective position effect. First, breakage of gene spans and enhancer-promoter interactions can cause mRNA truncation, chimeric transcripts, or misregulation of genes overlapping and near to breakpoints (Castermans et al 2007; Frisher et al 1986; Lupianez et al 2016; Ren and Dixon 2015). Previous work has found that common inversions in *D. melanogaster* are less likely to interrupt genic coding sequences that we expect under a random breakpoint model (Corbett-Detig and Hartl 2012), consistent with a possible role for natural selection in purging arrangements that interrupt genic sequences. Similarly, another work found that none of 54 polymorphic inversions in *D. pseudoobscura* interrupted a protein-coding gene span with their breakpoints (Fuller et al 2017). However, a mutational bias that preferentially creates breakpoints in intergenic regions is also consistent with these findings, making a comparison between rare and common inversion breakpoints a powerful framework for distinguishing among possible causes.

Second, factors related to local regulation, including polytene Topologically Associated Domain (TAD) boundaries, chromatin state, and insulator elements, might also have large impacts on the intrinsic fitness of newly formed arrangements. TAD boundaries generally demarcate related regulatory regions and disruption of these regions may alter local gene expression (Sexton et al 2012). Chromatin markers often determine local expression and repressive chromatin has been shown to spread across inversion breakpoints, repressing nearby gene activity in a process called position-effect variegation (PEV; Cryderman et al 1998). Insulator elements generally denote epigenomic boundaries between adjacent chromosomal regions and they can prevent the spread of heterochromatin via PEV and ectopic enhancer activity across insulator-maintained regions (Bushey et al 2008; Gaszner and Felsenfeld 2006; Sigrist and Pirrotta 1997; Yang and Corces 2012). High-frequency inversions should generally avoid disrupting local gene regulation through the manipulation of chromatin states or spatial compartmentation. Similarly high-frequency inversion breakpoints should colocalize with insulator elements, thereby mitigating regulatory impacts of large-scale rearrangements. In fact, inversions that have fixed between lineages show a strong association with insulator elements (Negre et al 2010), suggesting that we might observe similar effects within polymorphic high fitness arrangements.

Although suggestive, biased breakpoint distributions observed in high frequency and fixed arrangements are consistent with either natural selection or a mutational bias. As an example, let us suppose regions that border TADs are more likely to break due to physical stress or environmental exposure. We would expect to observe increased breaks in these regions not associated with the action of natural selection but instead resulting from a biased mutational process based on genomic context. Identifying and teasing apart these features therefore requires comparisons of inversions that have been exposed to natural selection to varying degrees, allowing us to draw an association directly between level of selection and qualities of the breakpoint region.

Here we evaluate these and other factors that may influence the intrinsic fitness of chromosomal inversions. We leverage population resequencing datasets from more than 1,000 *D. melanogaster* isolates to detect and *de novo* assemble both breakpoints of 18 rare naturally occurring inversions. We compare these “rare” inversions to known high frequency inversions in *D. melanogaster* (Corbett-Detig and Hartl 2012) as well as a set of inversions that have fixed between species in the *Melanogaster* subgroup (Ranz et al. 2007). For each inversion frequency category we compared their breakpoints to a permuted random breakage model and to every other category. In this framework, we expect mutational biases to appear as a difference in all comparisons with random expectations but not necessarily show an association with population frequency category, while features that influence fitness should both be different from random expectation and show an association with population frequency. By comparing rare, common and fixed inversion breakpoints, we uncover evidence that selection-driven position effects play an important role in shaping the fine-scale distribution of inversion breakpoints in natural populations.

## Methods

### Genome Version, Insulator, Chromatin, TAD and Gene Annotations

All our analyses are based on alignments to *D. melanogaster* genome version 6.26 (Hoskins et al 2015). We obtained genome annotation data including gene spans and mRNA coding regions from flybase. We obtained insulator binding site positions from (Negre et al 2010, accession GSE16245). TAD domains we present in this work are derived from polytene chromosomes (Eagen et al 2015) as well as additional TADs boundaries from specific cell lines (Ulianov et al 2016). As necessary, we converted the coordinates of genomic features from genome version 5 to 6 using the flybase coordinate batch conversion tool (https://flybase.org/convert/coordinates).

### Chromatin Data Source, Structure, Terms

Chromatin marker data was sourced from (Kharchenko et al 2011, accession GSE25321). We categorized regions of the genome by chromatin content into active, inactive, or mixed using data from Kharchenko’s nine-state model (Kharchenko et al 2011). Chromatin types were grouped by general activity as delineated in Kharchenko et al. (2011) into active chromatin and inactive chromatin. A break region was considered active or inactive if it exhibited only or primarily (70% or more of region) active or inactive chromatin markers and mixed if it exhibited similarly high levels of both. We investigated regions of up to 10kb to either side of each predicted breakpoint position for this analysis. We were particularly interested in examining inversions with breaks in different chromatin landscapes, as previous studies of “position-effect variegation” (PEV) indicated that inversions bringing genes into contact with heterochromatin can result in the suppression of those genes even if the spans themselves are undisturbed (Cryderman et al 1998; Eissenberg et al 1992; Puig et al 2004; Shatskikh et al 2018; Vogel et al 2008). We defined for our purposes a blending (and potentially PEV-inducing) inversion to be one that had breaks in different categories of chromatin, while referring to one with similar chromatin landscapes at either end as pure.

### Short Read Alignment

We obtained short read data as fastq files from the Sequence Read Archive. All sort read data is described in Lack et al 2016 and was originally produced in (Pool et al 2012, Lack et al 2015, Mackay et al 2012, Kao et al 2015). We aligned the short read data using bwa v0.7.15 using the “mem” function and default parameters (Li et al 2013). All post-processing (sorting, conversion to BAM format, and filtering) was performed in SAMtools v1.3.1 (Li et al 2009). We filtered these BAM files to include only those alignments with a minimum mapping quality of 20 or more.

### Breakpoint Identification

As in previous works that characterized structural variation using short-insert paired-end Illumina libraries (*e.g.* (CorbettDetig et al 2012, Cridland et al 2010, Rogers et al 2014), we first identified aberrantly-mapped read “clusters”. Briefly, here, a cluster is defined as 3 or more read pairs that align in the same orientation (for inversions, this is either both forward-mapping or both reverse-mapping) and for which all reads at one edge of the cluster map to within 1 Kb of all other reads in the cluster. We considered only aberrant clusters where both ends mapped to the same chromosome arm as the vast majority of inversions in *Drosophila* are paracentric (Krimbas and Powell 1992). We required that each cluster be a minimum of 500 Kb apart. We then retained only those potential inversions for which we recovered both forward and reverse mapping clusters there were within 100 Kb of one another. The choice of a maximum distance between possible breakpoint coordinates was included to reduce the possible rates of false positives and because none of the known inversions whose breakpoints have previously been characterized included a duplicated region of 100 Kb or more (Corbett-Detig et al 2012; Ranz et al. 2007). We further filtered all breakpoint clusters that overlapped annotation transposable elements as these are the primary source of aberrantly-mapping read clusters in previous works (*e.g.*, Corbett-Detig et al. 2012).

As an additional check for the accuracy of our newly discovered breakpoints, we compared our distribution of rare breakpoints to the known cytogenetic distribution and found no chromosomal or by-region differences (p=0.7, chi-square; cytogenetic data from Corbett et al 2016 who summarized Krimbas and Powell 1992). The short insert size from previous sequencing experiments ranged from ∼400bp to ∼800bp, which may have lead to a non-trivial false negative rate of cluster discovery particularly if the breakpoints contain repetitive elements or other large DNA insertions However, we don’t expect that these potential false negatives will bias our downstream analyses, and all previously characterized inversion breakpoints in the *Melanogaster* species complex occurred in unique sequence (Ranz et al. 2007, Corbett-Detig et al. 2012). All software used to perform these analyses is available from the github repositories associated with this project. Specifically, scripts used for breakpoint detection and assembly are in https://github.com/dliang5/breakpoint-assembly.

### Overlapping Inversions and In(2R)Mal

We also attempted to find sets of overlapping inversions. Briefly, for overlapping inversions, where one inversion arises on a background that contains another inversion, the breakpoint-spanning read clusters should be largely the same as inversions that arose on a standard arrangement chromosome. However, the key difference is that rather than pairs of forward and reverse mapping clusters, we expect to observe two distantly mapping read clusters in the reverse-forward and forward-reverse arrangements. We applied this approach for the 17 rare inversions that we initially discovered as well as to all samples that contained common inversions that are known from previous work (Corbett-Detig and Hartl 2012; Lack et al. 2015). We found only one such overlapping rare inversion, which is consistent with the known segregation distorter associated chromosomal inversion *In(2R)Mal* that is comprised of two overlapping inversions (Presgraves et al 2009). In our analysis here, we treat these overlapping inversions as independent, but our results are qualitatively unaffected if we simply exclude the second inversion.

### De novo Breakpoint Assembly

For each putative inversion, we then extracted all reads for which either pair mapped to within 5 Kb of the predicted breakpoint position. We converted all fastq read files to fasta and qual files as is required by Phrap, and we assembled each using otherwise default parameters but including the “-vector_bound 0 -forcelevel 10” command line options (Corbett Detig and Hartl 2012; Rogers et al 2014). We then used blast to align the resulting *de novo* assembled contigs to the *D. melanogaster* reference genome to identify the contig that overlapped the predicted breakpoint using the flybase blast tool (www.flybase.org/blast). We retained only inversions for which we could *de novo* assemble contigs overlapping both breakpoints, and we further discarded any contigs where the sequence intervening two distant genomic regions contained sequence with homology to transposable elements. All of the assembled breakpoint sequences are available in File S1. Assembly scripts are available from https://github.com/dliang5/breakpoint-assembly.

### Common and Fixed Breakpoint Positions

We obtained the breakpoint positions of common and fixed inversions from previous work. Specifically, common inversion breakpoints were reported in (Wesley and Eanes 1994, Andolfatto et al. 1999, Corbett-Detig and Hartl 2012, Lack et al. 2015). Fixed inversion breakpoints for the D. melanogaster subgroup were reported previously in Ranz et al. (2007). We converted coordinates for each from release 5 to release 6 as necessary using the flybase batch conversion tool (https://flybase.org/convert/coordinates).

### Defining Inversion Categories

There are three classes of inversion population frequency delineated in these analyses. Classically, the literature has referred to four categories of inversion, “common cosmopolitan”, “rare cosmopolitan”, “recurrent endemic”, and “unique endemic” (Mettler et al 1977; Krimbas and Powell 1992). The latter half of each of these terms refers to the geographic distribution of the inversion, which is not important for this particular analysis. As we are examining factors which can allow an inversion to achieve high frequencies, as long as an inversion has gone to high frequency in any population it should fall into our category “common”, while “rare” refers to inversions which were found in only single isolates with the exception of In(2R)Mal. Specifically, “common cosmopolitan”, “rare cosmopolitan”, and “recurrent endemic” will all fall under our label “common” while we refer to “unique endemic” as “rare” inversions, similarly to the analysis in (Corbett-Detig 2016).

The third class, “fixed” inversions, are inversions that have gone to fixation within one lineage during divergence of the *Drosophila melanogaster* subgroup and whose breakpoints have been characterized previously (Ranz et al 2007). Originally all fixed inversions occurred as unique events in a *Drosophila* ancestor. They subsequently spread until they reached fixation in populations ancestral to contemporary species in the *melanogaster* subgroup. These fixed inversions were discovered by comparing synteny block orientation between *D. melanogaster* and its relatives (Lemeunier and Ashburner 1976) and have been molecularly characterized previously (Ranz et al 2007). It is important to note that the vast majority of these fixed inversions occurred on the *Drosophila yakuba* branch and not in a direct *melanogaster* ancestor (Krimbas and Powell 1992; Ranz et al 2007). *Drosophila melanogaster* should therefore generally reflect the ancestral state and the genetic background on which these inversions originated rather than a derived state evolved after fixation. Common and rare inversions annotated here occurred in contemporary *D. melanogaster* and thus on a similar genetic background to that on which the *D. yakuba* inversions fixed. The functional annotations used here are also based on the *D. melanogaster* standard arrangement, meaning that all three frequency types should be comparable and have been selected in the context of the annotated functional landscape.

### Permutations and Statistical Tests

To compare inversion breakpoint positions to a randomized distribution, permutations for all categories of inversions (rare, common, and fixed) were performed with 1000 iterations of a group of randomly located breakpoints, holding the inversion number, lengths, and chromosome arms constant. Specifically, for each inversion, one thousand random points were chosen between the start of that chromosome arm and the end minus the length of the inversion-that is, from the entire set of possible starting points for an inversion of that size. Each randomly chosen point was used as the starting breakpoint for a simulated inversion, with the ending set at a distance further down the chromosome equal to the length of the real inversion in basepairs. Qualities of the genome at each of these breakpoints were recorded as our expected value for the random distribution of breakpoints. These tests were therefore designed to control for as many of the possible confounding factors as is feasible for this type of analysis, including bias related to the length and chromosome arm distribution.

All factors were tested for each frequency category in a pairwise fashion, as well as between each frequency category and an appropriate permuted set. We generally expect mutational biases to show association in all frequency categories versus permuted expectations but to be relatively invariant between categories in the absence of additional effects of natural selection. Factors that influence fitness, by contrast, should show no difference from expectation in rare inversions but should show increasing association in common and fixed inversion categories. More complex patterns involving both mutational biases and the effects of natural selection are also possible.

Tests were divided by the relationship with the factor. For factors that are a discrete numerical value for each break, such as distance to an element or length of a duplication, one-tailed Mann-Whitney U tests were used to compare the real distribution to a larger one thousand permuted set, yielding a single p-value per frequency category. Tests between categories of the distance-based factors and the duplication length test were performed distribution to distribution with pairwise Mann-Whitney U tests.

The second type of factor is a categorical value, such as interrupting a gene span or not. Rates of interruption were calculated for one thousand permutations. The percentile of the real rate of interruption was calculated to yield a single p-value for each distribution. All scripts used to produce the results of the permutation tests described above are available from the github repository associated with this project https://github.com/jmcbroome/breakpoint_analysis.

## Results and Discussion

### Inversion Breakpoints Discovered

We realigned all sequence data from over 1,000 *D. melanogaster* natural isolates that have been sequenced previously using paired-end sequencing methods (Grenier 2015; Langley 2012; Pool et al 2012; Mackay et al 2012; Lack et al 2015; Kao et al 2015), summarized in detail in Lack et al 2016. We identified 5,318 short read clusters that corresponded to possible inversion breakpoints that are a minimum of one Megabase from each other and for which we found both forward and reverse mapping read clusters (Figure S1). That is, for a given inversion relative to the reference genome, we expect to find a cluster of read pairs where both maps in the “forward” orientation and another cluster where each pair of reads both map in the “reverse” orientation (Corbett-Detig et al. 2012, see Methods). We also searched for overlapping inversions using a slight modification of this approach (Methods). To be as conservative as possible with our analysis, we retained only the set for which we recovered and successfully *de novo* assembled both breakpoints for a given inversion. Additionally we removed any putative breakpoint spanning contig that mapped with high confidence to multiple locations in the *D. melanogaster* reference genome. We ultimately retained 18 rare inversions. Three of our candidate rare inversions are corroborated by previous cytological evidence (Huang et al 2014; Presgraves et al 2009). Similarly, previous molecular evidence (Grenier et al 2015) similarly supports the identified breakpoints of three chromosomal inversions. The breakpoints of our putative rare inversions do not show unusual genetic distances from other samples isolated from the same populations, suggesting that these are relatively recent events and not older inversions that have undergone significant selection(Table S1).

The genomic and population distributions of rare inversions are largely consistent with our expectations based on extensive cytological work. First, our estimated rate of occurrence of rare inversions, 1.6% per genome, is within the range of estimates from cytological data across diverse populations 0.47%-2.71% (Aulard et al 2002; Krimbas et al 1992). Furthermore, we found no rare inversions on the X chromosome, which contains very few chromosomal inversions in natural populations of this species (Aulard et al 2002; Krimbas et al 1992). However, because we conservatively required that both breakpoints are detected from discordant short-read alignments and completely assembled *de novo*, and because we excluded any breakpoints that contained homology to annotated transposable elements, it is possible that our approach has underestimated the prevalence of rare inversions in these datasets.

### Disruptive Position Effects

#### Interactions with Gene Coding Spans and TADs

Inversions could be highly disruptive to sequences at their breakpoints. This has multiple classes of potential negative consequences, including the truncation of gene spans and the creation or alteration of ectopic enhancer interactions (Castermans et al 2007; Frisher et al 1986; Lupianez et al 2016; Ren and Dixon 2015). We investigated interactions with gene spans and regulatory marks under the hypothesis that higher-frequency inversions are less likely to cause large-scale disruptions to local functional elements. For each category, we calculated the percentile value of the real count of interrupting breakpoints against the permuted distribution, where low percentiles correspond to less breakage than expected. We found support for the avoidance of annotated mRNA coding spans among fixed (p<0.001, permutation) and common (p=0.015, permutation) inversions. Fixed inversions additionally exhibited avoidance of overall annotated gene spans (p<0.001, permutation) while common showed a similar trend (p=0.045, permutation). Rare inversions in contrast to both of these categories exhibited weaker avoidance (mRNA p=0.023; gene p=0.0125; permutation).

Because the proportion of gene-breaking inversions is inversely correlated to population frequency (44% of rare inversion breakpoints, 38% of common inversion breakpoints, 25% of fixed inversion breakpoints), our results are consistent with gene interruption being a selective position effect. We note here that the baseline rate of interruption is still relatively high even in the most conservative category, at 25% of fixed inversion breakpoints. This is likely due to the gene density of the *Drosophila melanogaster* genome and indicates that in at least some cases the interruption of a gene span may be effectively neutral or even positively selected. The trend across categories however indicates that there is at least an association of decreasing population frequency with gene interruption. We also note that rare inversions appear to interrupt genic regions less often than expected by chance. This particular point could be explained by the critical nature of some genes to survival; specifically, in that it must not have killed the living embryo we discovered it in. The preservation of gene spans by rare inversions may also be explained by a mutational bias of chromatin state or basepair composition favoring intergenic regions. As a final potential explanation we note that because many of the samples used in this work were inbred, either intentionally or passively as isofemale lines, inversions that induce recessive deleterious fitness effects might still be exposed to selection and purged from the line prior to sequencing.

Topologically associated domains, or TADs, are short three-dimensional structures representing spatial knots in the genome, often highly conserved and associated with coordinated gene regulatory blocks (Cavalli and Misteli 2013). We therefore expect that inversion breakpoints will interrupt TADs less frequently than under random breakage models if natural selection purges inversions that affect these large-scale regulatory features. Consistent with this, fixed inversion breakpoints exhibited avoidance of polytene TAD spans (p<0.001, permutation). Common and rare polymorphic inversions do not exhibit significant avoidance of TAD spans, however (rare p=0.1405; common p=0.111; permutation), though they both trend in the expected direction. We speculate that one potential explanation for this observation is that as *Drosophila melanogaster* are somatically pairing organisms with trans-homolog regulation (Fukaya and Levine 2017; Lewis 1954). The compartmentation of the inversion-free paired homolog may help rescue ectopic regulation in heterozygotes and enable inversions to persist at polymorphic frequencies if negative regulatory effects associated with interrupting TAD boundaries are largely recessive. Further exploration of the spatial conformation of the genome directly around these breakpoint regions in homo-versus heterozygotes is needed before stronger conclusions about this relationship can be made.

#### Sizes of Inverted Duplications

Paired staggered double-strand breakage is the major mechanism by which inversions events occur in the *D. melanogaster* subgroup (Puerma et al 2016; Ranz et al 2007). These breaks leave an overhang of sequence at each end of the putative inversion. After repair in inverted orientation, the result is inverted duplicated regions on either side of a new inversion with length equal to the overhang left over after the double strand break (figure 2A). A study in *Anopheles gambiae* has shown that the inversion *2L+*^*a*^ is viable presumably because it generates functional copies of disrupted genes (Sharakhov et al 2006). To disrupt a given functional span at the sequence level without creating a complete duplicate, both sides of a double-strand break must fall into that same functional span (figure 2A). Longer duplications are thus less likely to disrupt individual functional elements. Therefore we hypothesized that selection will favor longer duplicated regions that minimize impacts on local sequence function.

**Figure 1:**
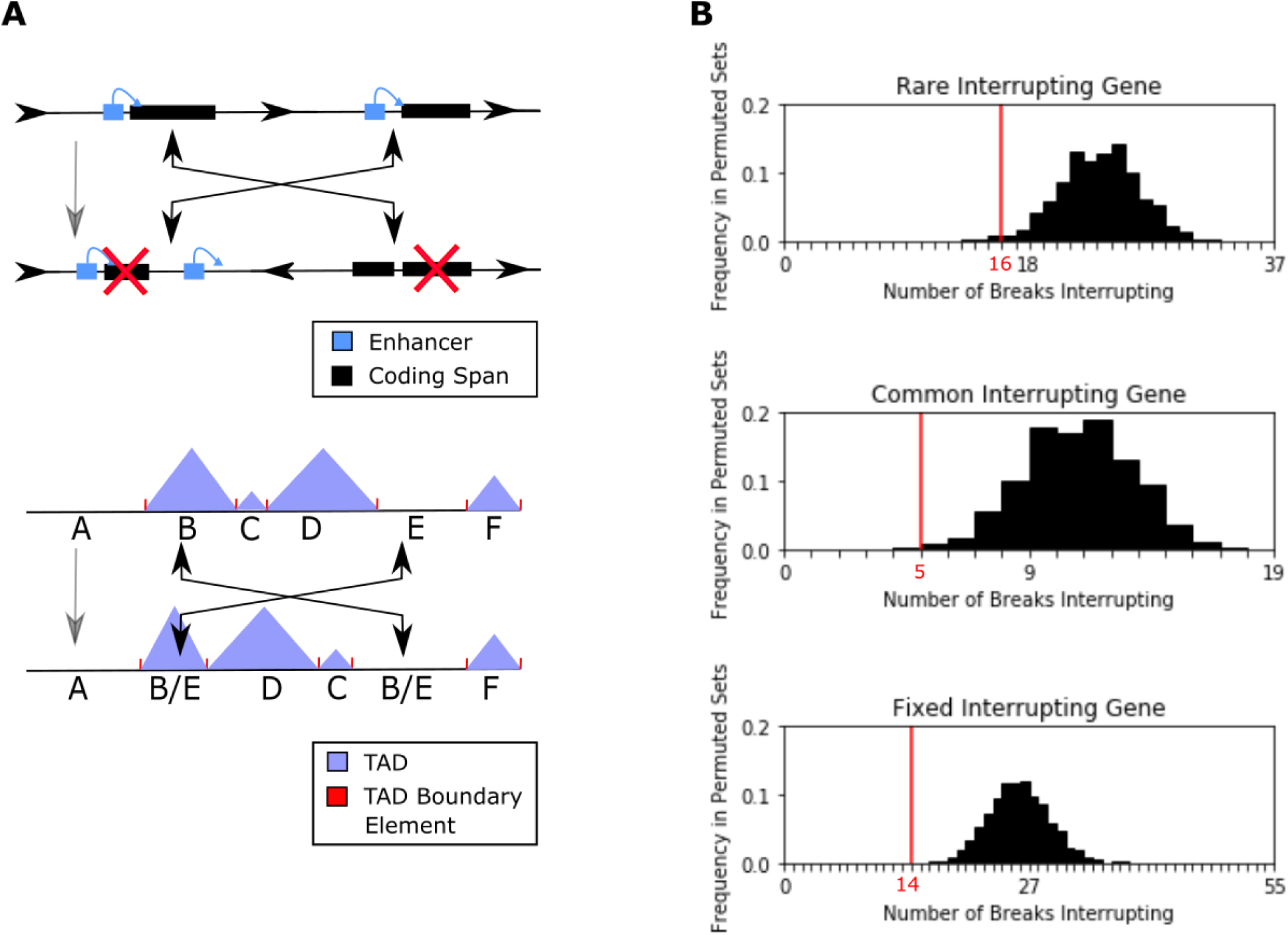
**A) Inversions can interrupt coding spans, disconnect enhancer/promoter pairs, and resize and remake TADs.** The top depicts a gene span being broken resulting in truncation (left) and an enhancer being separated from its target leading to silencing (right). The bottom depicts how topologically associated domains can have their gene content altered and size and order changed by inversion events. **B) All three breakpoint categories have a significantly lower rate of interruption of gene spans compared to a random breakage model**. In each plot, the x-axis represents the number of breaks interrupting genes. The histogram represents the number of genic interruptions for inversion breakpoint locations under a random breakage model with one thousand permutations. The actual value is represented by the vertical line in red.

**Figure 2:**
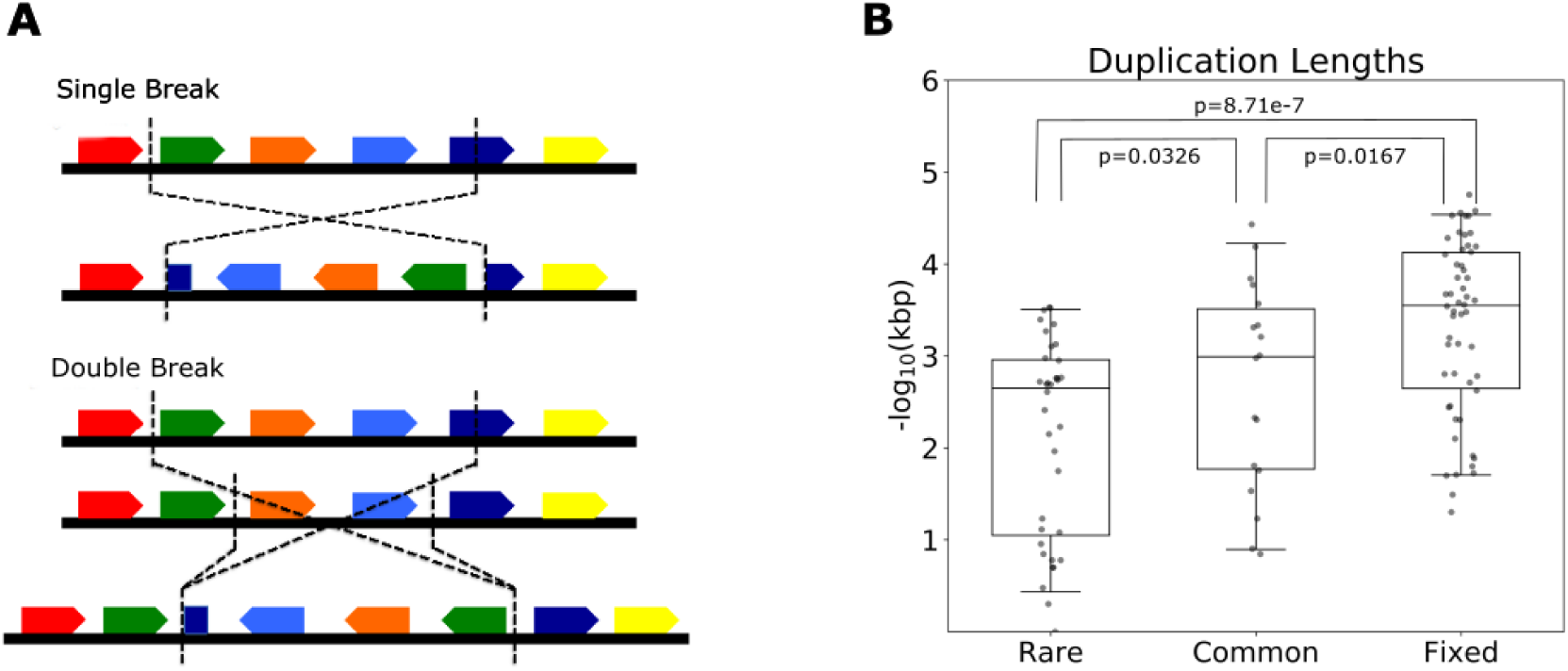
**A) Staggered breakpoints generate duplications that might suppress the impacts of sequence disruption.** This model figure shows the difference in effect of single and double break mechanisms. In the single break, the dark blue gene span is divided in half in the inverted line with no functional copies remaining. In the double break, the dark blue gene span is duplicated and a functional copy remains to the right of the break region. **B) Higher population frequency categories exhibit longer duplications.** The box and jitterplots represent the duplication lengths of each inversion class.

We found that inversions that have gone to fixation have significantly longer duplications than either common (p=0.01674, Mann-Whitney) or rare (p=8.717e-7, Mann-Whitney) inversions. Common inversions in turn have significantly longer breakpoint-adjacent duplications than rare inversions (p=0.03257, Mann-Whitney). Therefore, long duplication lengths are associated with increasing population frequency, which supports our hypothesis of long duplications acting as a compensatory mechanism for otherwise negative position effect by preserving intact functional elements or the proximal relationships among functional elements within duplicated regions. Formally, our analysis is consistent with higher relative fitness of inversions with longer inverted repeats, but does not necessarily require deleterious effects at single breakpoints (Figure 2A). It is also possible that inversions are positively selected when they contain larger breakpoint-adjacent duplications because of positive effects associated with gene duplications or chimeric gene products (Puerma et al 2016). However, given that microsynteny is largely maintained over evolution and given that the *Drosophila* genome contains a high density of functional elements, we favor our hypothesis that larger repeats are favored because they avoid disrupting functional elements. Nonetheless, positive fitness effects might also contribute to this pattern.

### Impacts on Chromatin Domains

#### Interactions with Chromatin Landscapes

Inversions do not merely relocate sections of genome sequence; they also transform the epigenetic landscape surrounding their breakpoints, relocating protein binding sites and creating new boundaries between distinct chromatin structures. These alterations can have effects that reach into neighboring regions, altering gene activity nearby to the break. The most-studied effect of this type is position-effect variegation (PEV). PEV occurs when an inversion has breaks in heterochromatic and euchromatic regions and the heterochromatin spreads to its newly juxtaposed active neighbor, silencing it (Cryderman et al 1998; Eissenberg et al 1992; Puig et al 2004; Shatskikh et al 2018; Vogel et al 2008).

While the physical mechanism of heterochromatic spread have been studied in detail, its potential role in inversion evolution remains unclear. As a brief reminder of terms, mixed chromatin is spans of genome with both active and inactive chromatin markers, as opposed to active spans and inactive spans which are primarily one or the other. Blending inversions are inversions with breaks in two different types of chromatin landscape, such as active and inactive or mixed and active, as delineated in methods. Blending inversions bring alternative chromatin types into close contact and allow for position-effect variegation or other effects, while pure inversions do not. The system is graphically explained in Figure 3A.

**Figure 3:**
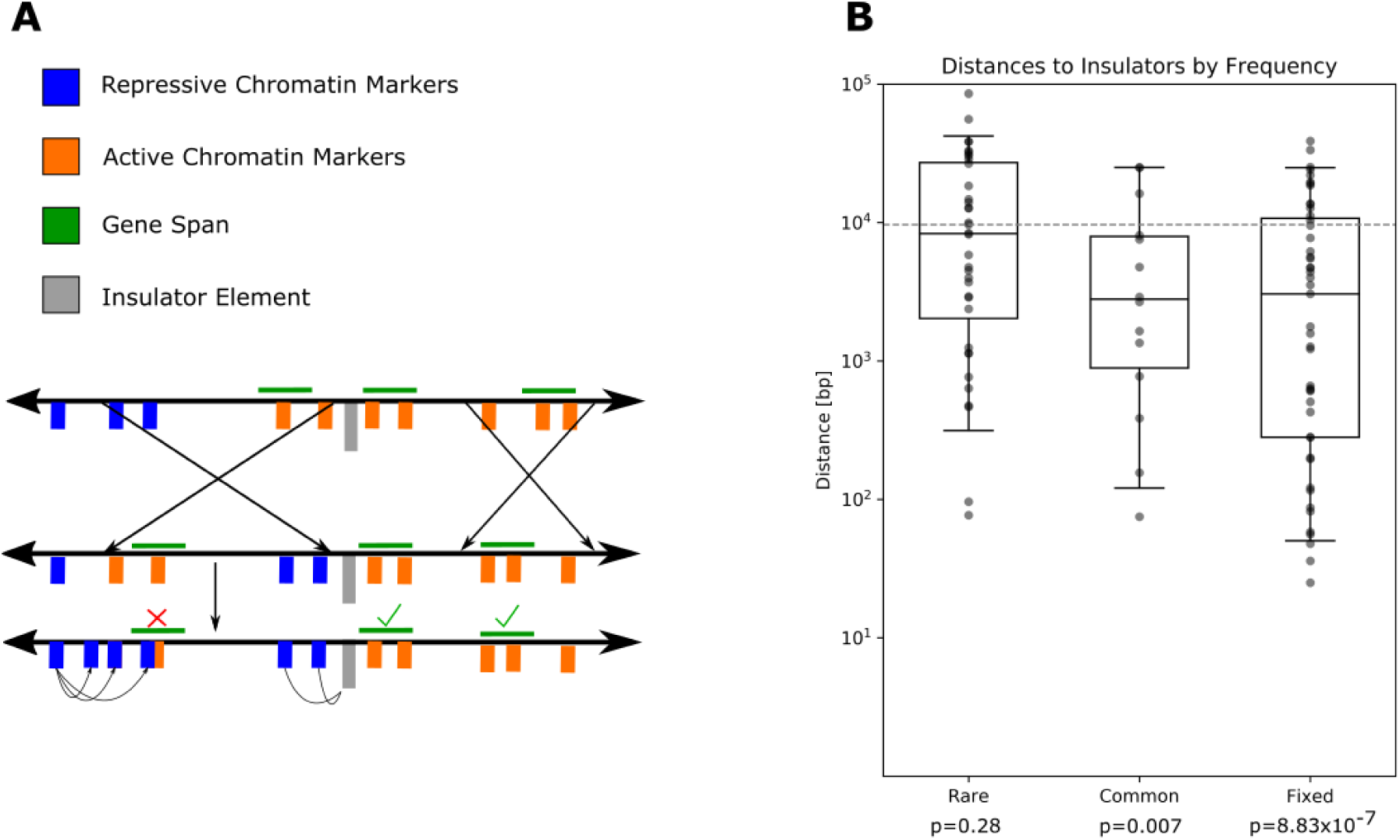
**A. Insulators prevent spread of repressive chromatin across blending breaks.** This pair of hypothetical inversions have four breakpoints that do not interrupt gene spans. The left inversion is a **blending** inversion, with one break in a repressive gene-free chromatin landscape indicated by blue chromatin marks, while the other is between two active genes in an active landscape. The right inversion is **pure** and has both breakpoints in active chromatin landscapes. An insulator element separates the two genes near the right breakpoint of the blending inversion. After inversion, one of the genes is relocated to be adjacent to repressive chromatin markers, which spread across the breakpoint through position-effect variegation and repress the gene. Position-effect variegation is blocked at the other blending inversion break by an insulator element, preventing any repressive effects. The pure inversion does not bring repressive and active chromatin into proximity and therefore position-effect variegation does not take place. **B. Higher population frequency correlates with insulator proximity.** The distributions of distances are from breakpoints to the nearest insulator element. Note that the y-axis is a logarithmic scale. The dashed grey line is the expected median distance of a random-breakpoint model. The p-values at the bottom are the Mann-Whitney U test result for being different from the permuted distribution (grey line).

We discovered an enrichment of fixed blending inversions (Figure S3A; p=1.014×10^−5^, Mann-Whitney). The age of the inversion may also play a role in this, as ancient fixed inversions may have evolved on a different chromatin distribution than that in either modern lineage. Insulators associated with these fixed inversions may also have served to form and maintain boundaries between shifting chromatin types over these breaks over evolutionary time. Nonetheless, this suggests that mixtures of distinct chromatin states is not a strong barrier to polymorphic inversion formation and fixation. We additionally found a weak tendency for rare inversion breakpoints to occur in constitutive heterochromatin (p=0.00228, permutation, figure S3A) without other biases (active chromatin p=0.296, mixed chromatin p=0.939, permutation, figure S3A). Common inversions did not exhibit any detectable pattern of breakage (active chromatin p=0.6, mixed chromatin p=0.438, inactive p=0.635, permutation, figure S3A). Fixed inversion breakpoints appeared random beyond the enrichment of mixed states discussed above (active chromatin p=0.538, inactive chromatin p=0.886, permutation, figure S3A). Our direct investigation of chromatin mixing did not find any significant differences in the rate of blending versus pure inversions for any group against each other or random expectation (Figure S3B). It is possible that mutational bias drives the occurrence of breakage in heterochromatic regions, but the relocation of heterochromatin into more active areas of the region and resulting PEV is somewhat deleterious, causing selection to effectively counteract the effects of mutational bias and result in a near-random pattern. The overall impact of PEV directly on polymorphic inversion location appears to be minimal. This may be due to weak selection against PEV silencing in nature or to compensatory mechanisms that prevent PEV from occurring in high-frequency inversions. The alteration of the chromatin landscapes, at least on the relatively broad scales we consider here, appears to play little role in shaping the fitness of newly arisen arrangements.

#### Interactions with Insulators

Insulator elements are key to the structure and function of genome regulatory networks, serving as structural anchor points, physical blockers of enhancer interactions, and boundary elements between TADs or chromatin compartments (Chung et al 1993; Negre 2010; Roseman et al 1993). Insulators may reduce or prevent PEV (Bushey et al 2008; Gaszner and Felsenfeld 2006; Sigrist and Pirrotta 1997; Yang and Corces 2012). Strong association with these elements may act as a compensatory mechanism that preserves chromatin states and thereby allows polymorphic inversions to occur at the boundaries between heterochromatic and euchromatic regions. Association with insulator elements may also prevent the ectopic interaction of enhancers between new neighbor gene networks. Previous studies have shown developmental disorders can occur in mammals from inversion rearrangements with no additional mutation via ectopic enhancer activity (Ren and Dixon 2015; Lupianez et al 2016). Notably, recent data suggests inversion breakpoints are rarely associated with local perturbed expression in *Drosophila* (Fuller et al 2016; Ghavi-Helm et al 2019; Said et al. 2018). However, there still appears to be some cases of ectopic enhancer/promoter interactions as in *subdued* and *Dscam4* (Ghavi-Helm et al 2019). This suggests that new ectopic interaction may be enabled by inversions but that these interactions rarely impact expression levels due to another mechanism.

Insulators can act as a compensator by blocking these interactions, thereby maintaining correct gene expression and enabling otherwise detrimental inversions to persist in natural populations (Chung et al 1993; Bushey et al 2008; Gaszner and Felsenfeld 2006; Roseman et al 1993; Wallace and Felsenfeld 2007; Yang and Corces 2012).These elements have already been previously shown to associate with fixed structural rearrangements that alter local synteny (Negre et al 2010). We hypothesized that the association with insulator elements may be an important factor mitigating fitness costs of chromosomal inversions and therefore would be correlated with population frequencies.

Our data show that common and fixed inversions differ substantially from expectations with respect to their median distances to insulator elements (rare p=0.28, common p=0.007, fixed p=8.83×10^-7, Mann-Whitney). We arranged the test against random expectations by generating a permuted distribution of insulator distance values and comparing the larger permuted distribution with the smaller real distribution of distance values of each category. Common inversion breakpoints are marginally significantly closer to insulators than are rare inversions breakpoints (p = 0.0589, Mann-Whitney) fixed and common inversion breakpoints are not statistically different (p=0.325, Mann-Whitney) and fixed inversion breakpoints are significantly closer to insulator elements than are rare inversions (p=0.0074, Mann-Whitney). We additionally searched for directional biases in insulator presence (i.e., whether insulators are found within the duplicated regions or outside) and found no evidence for any directionality to the association (Figure S2).

While these results are promising for insulators as compensation mechanism, the association is not directly proven here. A recent paper discovered a lack of gene expression differences over and around breakpoint regions (Ghavi-Helm et al 2019). We investigated insulators as a potential cause for this lack of misexpression on the balancer but found no enrichment (p=0.57, Mann-Whitney). Because they are maintained as transheterozygotes, balancer chromosomes are shielded from selection in homozygotes in a similar fashion to rare inversion and were artificially generated. It makes sense that we would not see a selective association with insulators for these reasons. The part that challenges our specific hypothesis is that Ghavi-Helm et al. found minimal differential gene expression around balancer inversion breaks despite the lack of insulator association. The association between insulator distances and population frequencies is robust in our data, but those results suggest it may not be because it prevents ectopic enhancer interaction. A potential explanation may be that insulators affect more subtle gene expression phenotypes than Ghavi-Helm et al studied. For example, if insulators decrease the variance in expression across cells, this might not be reflected in mean expression values obtained via bulk RNA-seq.

Insulator elements, chromatin state, and topological domain boundaries are all related aspects of spatial and temporal gene regulation (Cavalli and Mistelli 2013). It is curious that only insulator elements associate with inversion population frequency in our data, because in many species insulator elements are associated with topological domain boundaries and topological domains in turn usually show enrichment of either active or inactive chromatin markers across their span (Cavalli and Mistelli 2013). A potential resolution lies in the temporal nature of the data; both topological domains and chromatin state are unstable and change over the course of embryogenesis and cell differentiation (Jost et al 2014; Thomas et al 2011). By contrast, insulator binding sites are motifs identified in nucleotide sequences and are static throughout the organism’s life (Negre et al 2010). Since chromatin and topological domain data were collected from *Drosophila* embryos and embryo-derived cell lines, it may be that their configurations are specific to embryonic development. Static insulator binding sites are perhaps a better representation of adult spatial regulatory structures. Thus, the aggregate effect of an inversion on fitness may be better predicted by the proximity to insulator element binding sites.

## Conclusion

In this work, we present evidence that natural selection has played a role in shaping the fine-scale distribution of chromosomal inversion breakpoints in the *D. melanogaster* subgroup. Although it is possible that breakpoint effects sometimes increase the fitness of new arrangements, e.g. by creating new expression patterns of transposed loci or chimeric transcripts (Puerma et al 2016), our data support the conclusion that selection acts against breakpoints that interrupt functional elements within breakpoint regions to purge deleterious fitness consequences, consistent with studies in other *Drosophila* species (Fuller et al 2017). In particular, we found evidence that insulator elements are important for improving fitness of chromosomal inversions.

A stronger understanding of the factors underlying the distribution of polymorphic inversions is requisite in the study of their evolutionary impact. These factors determine what parts of the genome can have linkage preserved by polymorphic inversions to form complex phenotypes. Variations in these factors may also explain differences in the formation and role of inversions across species. Before an inversion can be selected for recombination suppression or other features it must form in a genomic context where breakage is not immediately and strongly detrimental. Given two additional features of chromosomal inversions, (1) the *de novo* mutation rate is likely very low (Krimbas and Powell 1992), and (2) the conditions for a new arrangement to be favored by natural selection are sometimes restrictive (Charlesworth et al 2018; Kirkpatrick et al 2006; Hoffmann et al 2008), the impacts of fine-scale inversion breakpoint positions on the fitness of new arrangements suggest that the availability of suitable, high fitness arrangements may often be rate-limiting for adaptive evolution when suppressed recombination is favorable.

## Supporting information

Supplementary Figures and Descriptions of Files

Supplemental Table 1 (Breakpoint divergence)

Supplemental File 1 (Breakpoint fasta)

## Acknowledgements

The authors give their thanks to Stephen Schaeffer and Zach Fuller for their detailed suggestions in manuscript preparation and Iskander Said for his assistance collecting the low-frequency inversion data. This work was supported by the National Institutes of Health (R35GM128932)

