## Supplementary Figures and Descriptions of Files for "Fine-scale position effects shape the distribution of inversion breakpoints in *Drosophila melanogaster*"

**Supplemental Materials**

**Explanation of Permutation Testing**

**Figure S1** Breakpoint confirmation.

**Figure S2** Bidirectionality of insulator association.

**Figure S3** Chromatin states over breakpoints

**Table S1** Rates of base mismatch/sequence divergence of all breakpoint regions

**File S1.** Fasta file containing all assembled breakpoint-spanning contigs.

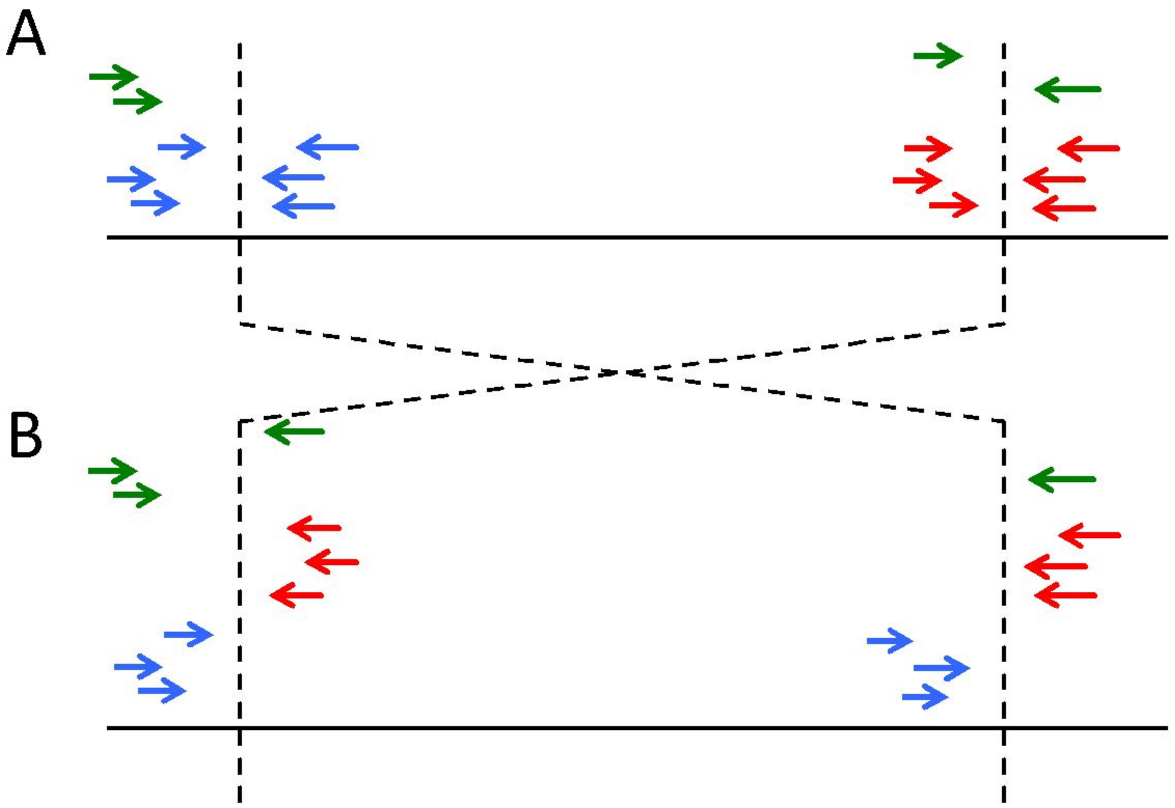


**Figure S1.** (A) Mapping positions along the genome from which short reads that support the presence of inversion breakpoints are derived. (B) Mapping positions along the genome where those reads will map on the standard arrangement reference genome that does not have this inversion. Read pairs that map in forward-forward orientation on the reference genome are shown in blue, read pairs that map in reverse-reverse orientation are shown in red. Finally, reads whose pair does not map, presumably because it overlaps the breakpoint, are shown in green. All reads show, as well as their unmapped pairs and additional adjacent standard mapping read pairs, were used to *de novo* assemble breakpoint adjacent regions.

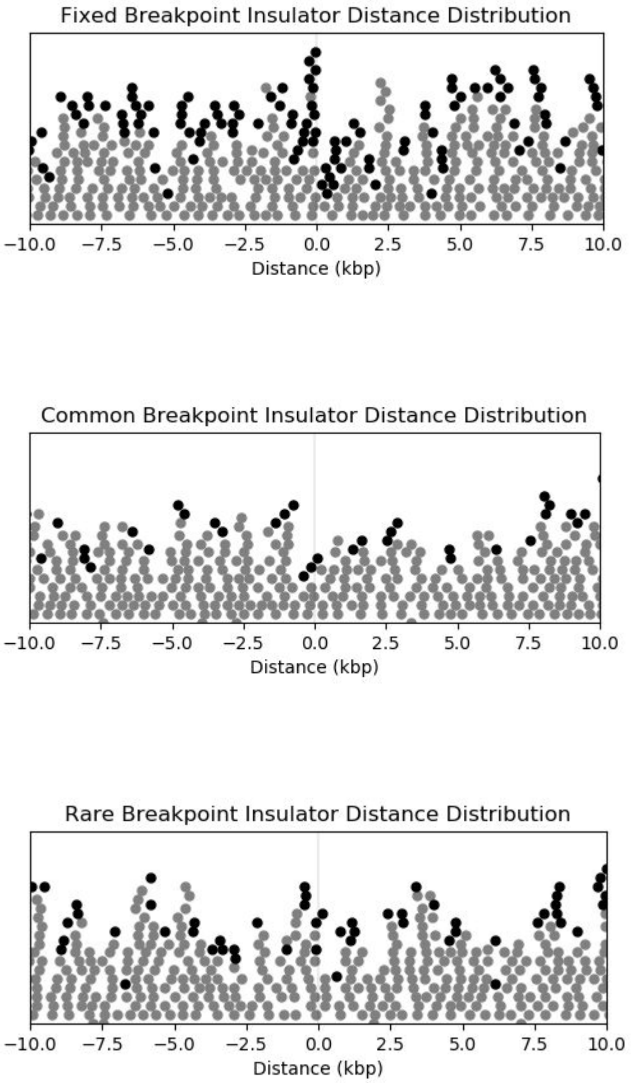


**Figure S2**: Vertical swarmplot of insulator distance values in a 10kbp window on either side of the set of breakpoints. Grey dots are permuted expectations; black is real data. No directional enrichment is evident.

A

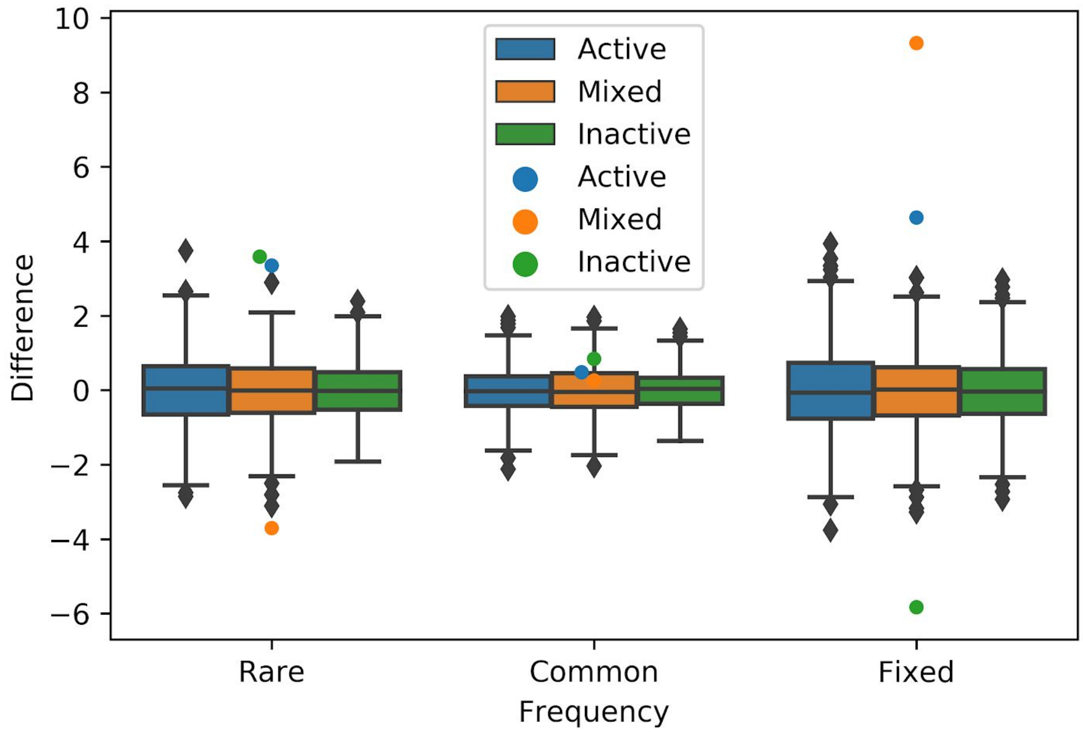


B

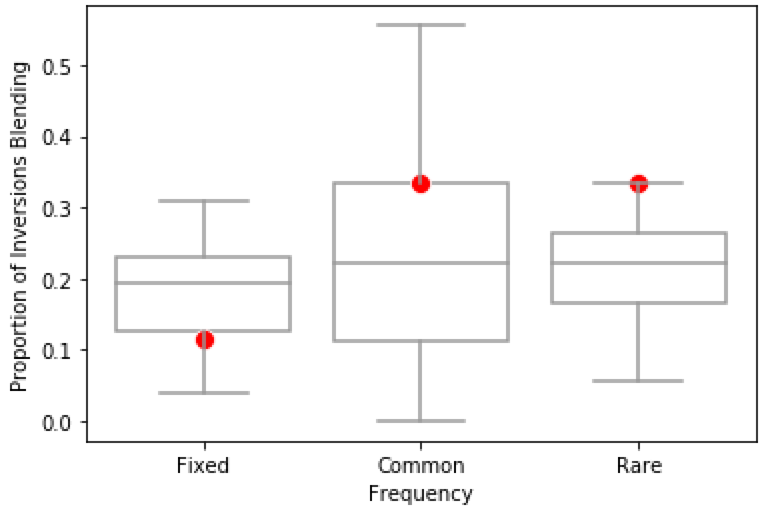


**Figure S3: A- Breakpoints are largely unbiased to chromatin states except towards mixed states in fixed inversions.**This series of boxplots with counts of inversion breakpoints per frequency category exhibits the three delineated chromatin activity states. Actual values are represented as points with the permuted expectation for each category as boxplots. The y-axis is the number of breaks of that type more or less than the expected value of a permuted null model. We see a significant increase in mixed chromatin states across fixed inversions. While there some pattern of rare inversions blending chromatin less, it does not appear to be statistically significant. **B- No frequency category exhibits significant bias towards or against blending chromatin landscapes across their breakpoints.** These boxplots are of the proportion of total inversions that are blended as red dots against the background expected distribution of blending rate. We see no category shows strong evidence of either increased or decreased occurrence of chromatin blending compared to permuted expectations, though we see a non-significant enrichment of blending inversions in rare frequencies.

**Table S1 Description**

This tab delimited file displays for each breakpoint the name and sequence divergence of the breakpoint as well as the most diverged strain and its value, least diverged strain and its value, and the number of stains missing assemblies over the breakpoint region. Divergence values were first calculated as "# of mismatches / (# of matches + # of mismatches)" between each pair of strains with an assembly, where a match was when two bases at a point were concordant and mismatch discordant, discounting any points where either assembly in the pairwise was ambiguous. Each strain then had its mean divergence value from the set of comparisons to all other strains calculated and the distribution of those values was used to calculate the percentile of average divergence. Some inverted strains have no assemblies over the predicted breakpoint, leading to an "N/A" under percentile and "AMBIG" under the divergence value. This table only includes versions for which an assembly of the chromosome arm for that strain was available. No annotated inversion-bearing strain was either the most or least divergent strain in its population in a window of 10 kilobases around the breakpoint, indicating that these inversions are likely not exceptionally old.
